# Asymmetric Fin Shape changes Swimming Dynamics of Ancient Marine Reptiles’ Soft Robophysical Models

**DOI:** 10.1101/2024.02.15.580532

**Authors:** Hadrien Sprumont, Federico Allione, Fabian Schwab, Bingcheng Wang, Claudio Mucingat, Ivan Lunati, Torsten Scheyer, Auke Ijspeert, Ardian Jusufi

## Abstract

Animals have evolved highly effective locomotion capabilities in terrestrial, aerial, and aquatic environments. Over life’s history, mass extinctions have wiped out unique animal species with specialized adaptations, leaving paleontologists to reconstruct their locomotion through fossil analysis. Despite advancements, little is known about how extinct megafauna, such as the Ichthyosauria one of the most successful lineages of marine reptiles, utilized their varied morphologies for swimming. Traditional robotics struggle to mimic extinct locomotion effectively, but the emerging soft robotics field offers a promising alternative to overcome this challenge. This paper aims to bridge this gap by studying *Mixosaurus* locomotion with soft robotics, combining material modeling and biomechanics in physical experimental validation. Combining a soft body with soft pneumatic actuators, the soft robotic platform described in this study investigates the correlation between asymmetrical fins and buoyancy by recreating the pitch torque generated by extinct swimming animals. We performed a comparative analysis of thrust and torque generated by *Carthorhyncus, Utatsusaurus, Mixosaurus, Guizhouichthyosaurus*, and *Ophthalmosaurus* tail fins in a flow tank. Experimental results suggest that the pitch torque on the torso generated by hypocercal fin shapes such as found in model systems of *Guizhouichthyosaurus, Mixosaurus* and *Utatsusaurus* produce distinct ventral body pitch effects able to mitigate the animal’s non-neutral buoyancy. This body pitch control effect is particularly pronounced in *Guizhouichthyosaurus*, which results suggest would have been able to generate high ventral pitch torque on the torso to compensate for its positive buoyancy. By contrast, homocercal fin shapes may not have been conducive for such buoyancy compensation, leaving torso pitch control to pectoral fins, for example. Across the range of the actuation frequencies of the caudal fins tested, resulted in oscillatory modes arising, which in turn can affect the for-aft thrust generated.

## Introduction

Ichthyosauriformes are one of the most successful lineages of marine animals living from the Mesozoic time period (between ca. -252 and -66 million years (Ma)) based on taxonomic diversity and clade longevity (Moon & Stubbs 2020). They existed from the Early Triassic (-250 Ma) to the early Late Cretaceous (-95 Ma). Ichthyosauriformes were almost globally distributed and adapted to various aquatic habitats and lifestyles, ranging from near-shore coastal dwellers to open ocean cruisers (Motani 2005, Bernard et al. 2010, Motani et al. 2018). The earliest representatives of the clade are species with elongated bodies and seam-like tail fins fitting with a general anguilliform swimming mode (e.g., *Chaohusaurus* from China, (Maisch 2001)). Sub-carangiform to carangiform modes became dominant in clades following the Middle Triassic (e.g., *Mixosaurus* and *Besanosaurus* from Switzerland, (Dal Sasso & Pinna 1996, Ji et al. 2016); *Guizhouichthyosaurus* from China, (Pan et al. 2006)), whereas thunniform modes are typical for most post-Triassic parvipelvian ichthyosaurs (e.g., *Temnodontosaurus, Stenopterygius* and *Ophthalmosaurus* from Europe (Buchholtz 2001, Motani 2005, Gutarra et al. 2019)).

Body-size evolution shows extremely large forms during the Middle (Cymbospondylidae) and Late Triassic (Shastasauridae) instead of a continuous increase in size from the Early Triassic to the early Late Cretaceous (Sander et al. 2021). The general evolutionary pattern observed in ichthyosaurs indicates a transition from a streamlined symmetrical tail to an asymmetrical lobed tail resembling that of sharks but reversed (in that the ventral lobe is elongated and supported by the vertebral column), and eventually to a more rigid symmetrical lunate tail (Crofts et al. 2019). This trend suggests that natural selection favored enhanced locomotor abilities for this group of marine animals. A similar pattern is also observed in both cartilaginous and bony fishes (Flammang 2010).

Symmetrical and asymmetrical caudal fins are commonly referred to as homocercal and heterocercal, respectively. Heterocercal is also used to designate more specifically the unequally-lobed fins where the dorsal lobe is larger than the ventral one; to avoid confusion, we will refer to this type of fin with the less-common term hypercercal fin. On the other hand, when the ventral lobe is larger than the dorsal one, the fin is referred to as hypocercal (Giammona 2021, Lauder 2000, Wright 1878). In general, heterocercal tails are thought to contribute to the balance of vertical forces as they swing. This hypothesis is supported by the greater representation of hypercercal fins in negatively buoyant sharks (Lauder 2000, Giammona 2021) and hypocercal fins in positively buoyant Ichthyosauriformes (McGowan 1992). Each need to compensate their imbalanced buoyancy with forces generated by active swimming.

In the study of swimming mechanics, robots are demonstrating an increasing capability to replace living subjects in targeted experiments (Matta et al. 2019, Van Buren et al. 2017, Low & Chong 2010). In contrast to computer simulations, robot-based physical models are vital because they interact with the real world and allow testing specific hypotheses pertaining to biomechanics and control (McInroe et al. 2016, Ijspeert et al. 2007, Tangorra et al. 2009, Aguilar et al. 2016). These robots can be used to study locomotion at a system level, where hardware (material, morphology) and software (sensors, control) must work together to mimic natural movements (Nyakatura et al. 2019, Thandiackal et al. 2021). Using robots also allows testing movements that animals are not able to do or that could even be harmful to them (Ijspeert 2014), allowing the exploration of capabilities and behaviors of extinct animals that are not directly observable.

Bio-inspired robots are becoming closer and closer to their natural counterparts. In Tytell et al. (2016), the authors use an anguilliform robotic fish platform made of eleven motors and a passive tail to study the effects of tail stiffness in the undulatory swimming motion. In McHenry et al. (1995), the authors compare the swimming performance of a muscle-stimulated pumpkinseed sunfish with its casted fish replicas with different body stiffness. Soft robotics, in particular, has helped bridge the gap between natural and artificial systems by creating softer and more compliant components than we see in most traditional robots (Jusufi et al. 2017). This enables the creation of bio-inspired designs that can passively perform more life-like motion (Suzumori et al. 2007, Wright et al. 2019).

This work proposes a novel instrumented soft robotic platform to validate hypotheses about ichthyosauriform locomotion, complementing previous studies based on collected data from extant animals combined with mathematical models. Motani (2002) studied the effects of caudal fin propulsion and the speed of ichthyosaurs, and Sánchez-Rodríguez et al. (2023) analyzed the effects of the tail beat frequency in underwater undulatory swimming. They achieved this goal by analyzing in depth the thrust production associated with the different caudal fin morphologies displayed throughout the evolution of Ichthyosauria and measuring the impact of the heterocercal caudal fins on produced forces. The results obtained in this work show a direct correlation between heterocercal fins and the pitch torque generated while swimming, validating the hypothesis that the oscillating motion of the tail while cruising could compensate for the not-neutral buoyancy of such animals.

## Materials and Methods

### Soft Active Material Physical Model

The soft robotic fish-shaped marine reptile used in this work is presented in Figure 2. It consists of a flexible foil, representing the backbone, to which two soft pneumatic actuators are glued to provide bending actuation (Wolf & Lauder 2021). A frontal 3D-printed ABS cuff serves as an attachment point for the force/torque (F/T) sensing equipment (ATI nano17), while another 3D-printed, flexible, TPU A95 tail cuff allows the attachment and detachment of caudal fins for tests with different shapes.

The main body is made following a similar method as described by Jusufi et al. (2017) and Schwab et al. (2024), and based on the work of Mosadegh et al. (2014). The flexible backbone foil (0.52 mm thick shim stock: Artus, Inc.) has a flexural stiffness comparable to that of a fish (9.9e-4 [Nm^2^]) (Jusufi et al. 2017), and the actuators, made using a silicone-based elastomer (Dragon Skin™ 20, Smooth-On Inc.), are glued using a dedicated silicone adhesive (SilPoxy™, Smooth-On Inc.). The layer of SilPoxy used to glue the actuators is thin enough not to significantly affect the bending capabilities of the soft robotic platform. Furthermore, by using the same platform, the bending stiffness is the same throughout all the experiments, allowing for relative comparisons among different fins.

Caudal Fin Morphology

The different ichthyosaur fin shapes compared in this study were chosen from a subset of the different genera presented by Gutarra et al. (2019). They were selected for their shape variety and the significant coverage of the evolution period: *Cartorhynchus* (-250 Ma, Early Triassic), *Utatsusaurus* (-250 Ma, Early Triassic), *Mixosaurus* (-245 Ma, Middle Triassic), *Guizhouichthyosaurus* (-235 Ma, Late Triassic), and *Ophthalmosaurus* (-155 Ma, Middle Jurassic-Early Cretaceous). Reconstructing the tail shape is not an easy task, and fins can have similar but not identical shapes, as in the case of the *Guizhouichthyosaurus*, which is shorter and with the lower lobe more ventrally angled in the reconstruction made by Gutarra et al. (2019) than in the one of Jiang et al. (2020). In the end, the more dramatically angled fin has been kept to provide an extreme hypocercal fin case in this study. The cutoff between the caudal fin and tail fin has been arbitrarily chosen at the narrowest point of the fin to have a better attachment to the actuated backbone of the robotic platform. As a consequence, although the reproduced fins do not cover exactly the same internal morphologies across the tested subjects, they still provide valuable information regarding torques and forces generated while swimming. The different fin shapes are displayed in Figure 1. An additional rectangular fin (also referred to as Truncated (Tru)) was also tested for comparative measurements with previous work of Jusufi et al. (2017).

**Fig. 1.**
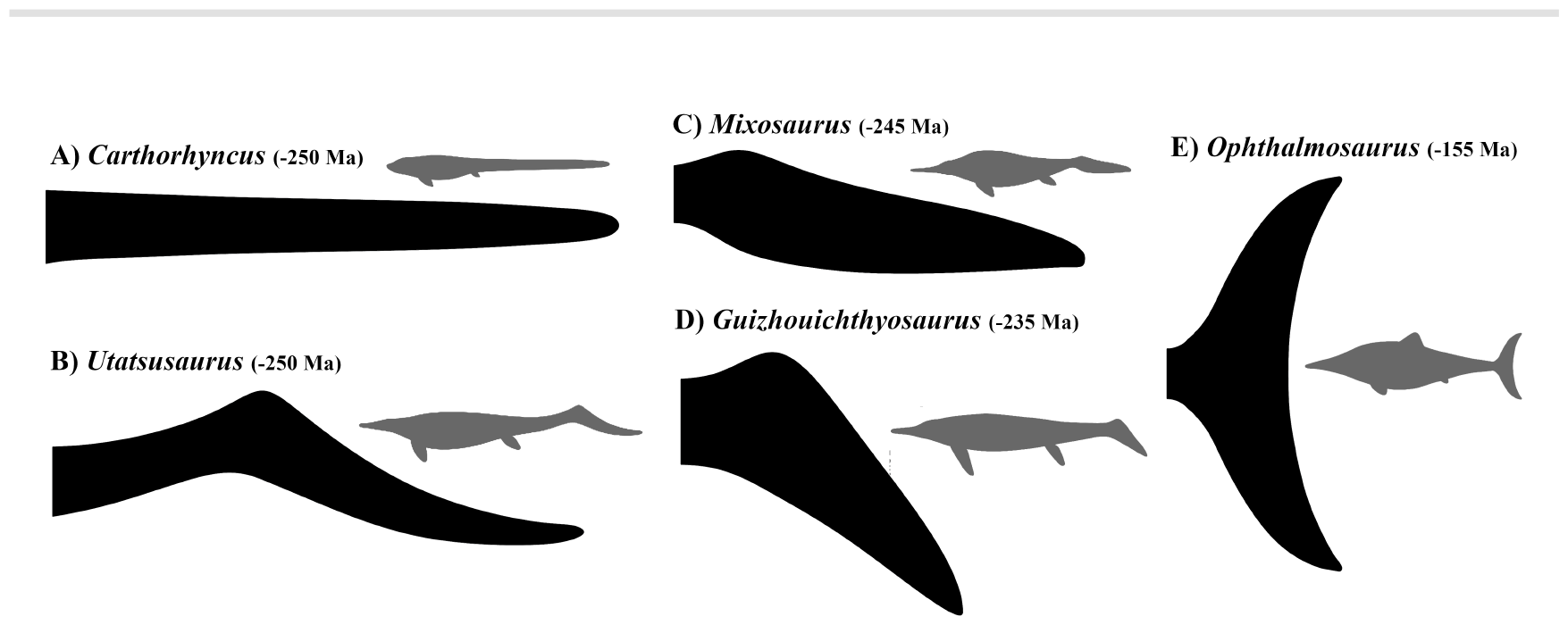
Heterocercal (B, C and D) and homocercal Caudal (A and E) fin shapes inspired by ichthyosaur model systems tested and compared in the flow tank experiments. Videos of the motion of five ichthyosaur-based caudal fins from a side view can be found in the supplementary material Movie 1. A) *Carthorhyncus*, B) *Utatsusaurus*, C) *Mixosaurus*, D) *Guizhouichthyosaurus*, E) *Ophthalmosaurus* tail fins

**Fig. 2.**
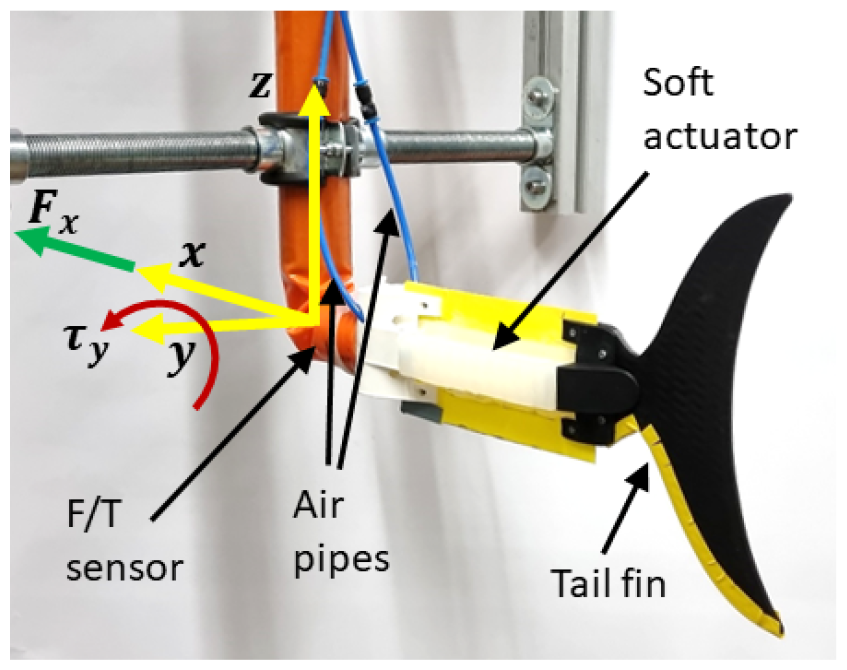
Soft robophysical platform utilized to measure pitch effects of hetero-versus homocercal fins in the flow tank experiment. *F*_*x*_ and *τ*_*y*_ represent the thrust force and the pitch torque along the *x* and *y* axis respectively. The Figure displays the *Ophthalmosaurus* fin shape.

The ichthyosaur fins are 3D-printed with flexible TPU A95 as flat foils with a thickness of 3mm, while the truncated is of the same material as the body to resemble Jusufi et al. (2017) and validate the testing apparatus. All the fins were scaled to present approximately the same surface area. Table 1 summarizes some shape parameters.

**Table 1.**
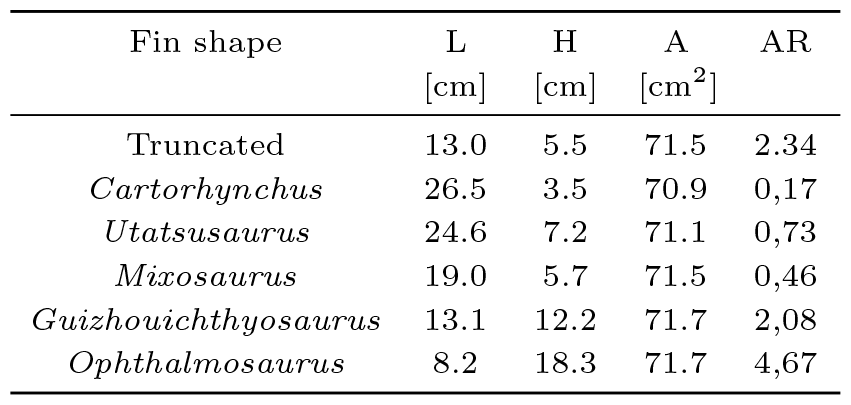
Geometrical properties of the 3D-printed tested caudal fins. L, H, A, and AR represent respectively length, height, surface area, and aspect ratio of the fin. AR is computed as AR = H^2^*/*A.

### Control System

The actuators are controlled via an off-board real-time microcontroller (myRIO, National Instrument) connected to a pressure reguator (ITV0050-3BS, SMC) and solenoid valves (SYJ7320-5LOU-01F-Q, SMC). The control loop frequency is 1 kHz. The signal to the valves controlling each actuator is a square wave that can be modulated in frequency and duty cycle (the percentage of the ratio of an active actuator to the total period of the waveform). The system pressure is maintained at a chosen value through the regulator by a Proportional Integral (PI) controller. The actuation pressure and the duty cycles are maintained constant across all experiments to ensure consistency in power output across different tails. The oscillating frequency range has been selected not to exceed the robot’s power capacity, as indicated by preliminary tests on amplitude reduction at higher frequencies. The robotic bending motion is otherwise performed open-loop. The soft robotic apparatus is then held by a pair of solid masts inside the recirculating flow tank at Empa Zürich. The flow tank has a 1*m ×* 0.6m cross-sectional area and can produce a controlled constant laminar flow around the apparatus of 0.02 *−* 1.5 m/s.

### Experimental Protocol

To measure the various fin performances, a 6-axis Force/Torque (F/T) sensor (ATI nano17), sampled at 1 kHz, is fixed to the front of the robotic fish-shaped marine reptile. It measures the thrust, side, and lift forces (F_*x*_, F_*y*_, and F_*z*_) as well as the roll, pitch, and yaw torques (T_*x*_, T_*y*_, and T_*z*_) produced by the platform while swimming. A high-speed camera with a frame rate of 400 Hz also records the body and fin motion from underneath to reconstruct offline the midline kinematics.

For every measurement conducted in this study, five distinct parameters were subjected to variation, namely actuation frequency, duty cycle, flow speed, and fin shape. The comparative analysis of each fin’s performance was undertaken via two comprehensive parameter sweeps. The first sweep involved the incremental adjustment of actuation frequency, while the second entailed systematic increments in water flow speed. The system pressure was maintained at 0.7 Bar throughout the experiments. Additionally, the duty cycle was intentionally set slightly above 50%, ensuring that both actuators operated concurrently for specific durations. This strategy was implemented to mitigate sudden directional changes in the fin’s motion, thereby averting potential F/T sensor saturation issues. Furthermore, as proved by Jusufi et al. (2017), a small amount of co-contraction leads to increased thrust generation. The distribution of the duty cycle was 0.55 for the long and slender fins (*Cartorhynchus, Utatsusaurus*, and *Mixosaurus*), 0.6 for the rectangular fin, and 0.65 for the shorter fins with a high aspect ratio (*Guizhouichthyosaurus, Ophthalmosaurus*). The higher co-contraction allows for a smoother transition between bending phases, avoiding abrupted changes of direction which would have caused the saturation of the F/T sensor.

The experimental procedure starts with the robot positioned in the water in an idle state. Subsequently, it initiates oscillatory movements, systematically augmenting the amplitude of these oscillations until reaching a state of constant amplitude, indicative of steady swimming under the same input command. Following this, the actuators are deactivated, and the robot returns to its initial idle state.

Each experimental trial involves the recording of data spanning a duration equivalent to 10 undulation periods of the tail during the phase of steady swimming. Notably, the transitional phases from idle to steady swimming and from steady swimming back to idle are excluded from the data collection. Both the initial and final idle segments are recorded, and their respective datasets are used to eliminate any inherent bias and to compensate for potential drift in the experimental observations.

The collected F/T data is passed through a median filter (50 samples window size) followed by a second-order Butterworth filter to remove the quantization noise with a cutoff frequency of 60 Hz. The steady swimming periods are then extracted to compute each axis’s mean forces and torques.

Fluctuation in the robot’s buoyancy occurs over the course of an actuation period as the actuators alternate between inflation and deflation during measurements. This dynamic behavior generates substantial bias forces on the F/T sensor that must be properly addressed. The buoyancy effect has been estimated by recording the F/T sensor’s data in static conditions initially with only one actuator inflated and then with both actuators. The forces in the x and z axes and the pitch torque measured this way are subtracted from the measured values during the experiments according to the duty cycle value, compensating for the buoyancy caused by the air in one or both actuators. Although negligible for the forward thrust, the buoyancy effect of the actuators has an impact on the measured values of the force in the z direction and the pitch torque, with peak values (measured when both actuators are inflated) respectively of 0.11 N and 2.5 Nmm.

### Data Availability

The datasets for this study can be found here https://github.com/ardianet1/finshape. Other data presented in this paper can be made available upon reasonable request.

## Results

The thrust data collected in this study is not an approximation of the actual animals’ overall swimming performance, as drag caused by the full body is not modelled, nor is the exact body-caudal motion of each specimen. Instead, the objective is to compare the effects of the various fin morphology changes while all other parameters remain untouched.

### Flow Speed Sweep

Figures 5, 6 and 7 show the average forward thrust (F_*x*_), vertical thrust (F_*z*_), and pitch torque (T_*y*_) during the flow speed sweep experiment.

The robot’s self-propelled swimming speed for a given combination of parameters is estimated by measuring the forward thrust at various flow speeds. Figure 5 shows a reduction of the forward thrust as the flow speed increases, similar to the results obtained by Wolf & Lauder (2021) in figure 8.4.C. The relation between flow speed and measured thrust is shown to be approximately linear, allowing the computation of the estimated self-propelled speed via linear regression. The fin motion produced by the platform allows the fins to produce thrust via added-mass. Lift-based swimming, which is exploited by thunniform swimmers like the *Guizhouichthyosaurus* and the *Ophthalmosaurus*, requires a specific motion pattern and fin profile that the current soft robotic platform does not provide. As such, it is expected that the longer fins generate more thrust in this specific experimental configuration than the short ones with higher aspect ratios, despite the latter usually being associated with high-speed swimming (Webb 1984, Sfakiotakis et al. 1999). The results are shown in Table 2. The *Utatsusaurus* displays the greatest self-propelled speed despite not being the longest fin. This result may indicate that the peculiar shape of the fin can help reduce drag. Indeed, as the robot swims, the distal section of the fin has a tendency to naturally flex upwards instead of staying aligned with the midline, thus presenting a reduced and angled cross-sectional area to the incoming flow. The *Guizhouichthyosau-rus* fin gets the worst result, which likely comes from the fact that those high aspect-ratio fins are usually associated with lift-based swimming, thus performing sub-optimally in this experiment. As the fasted cruisers today have a lunate or thunniform tail shape, such as great white sharks, tunas, or sword-fishes, the authors attribute the counter-intuitive low value for the *Ophthalmosaurus* to the lack of rigidity of the 3D printed fin and the non-thunniform tail swimming motion.

**Table 2.**
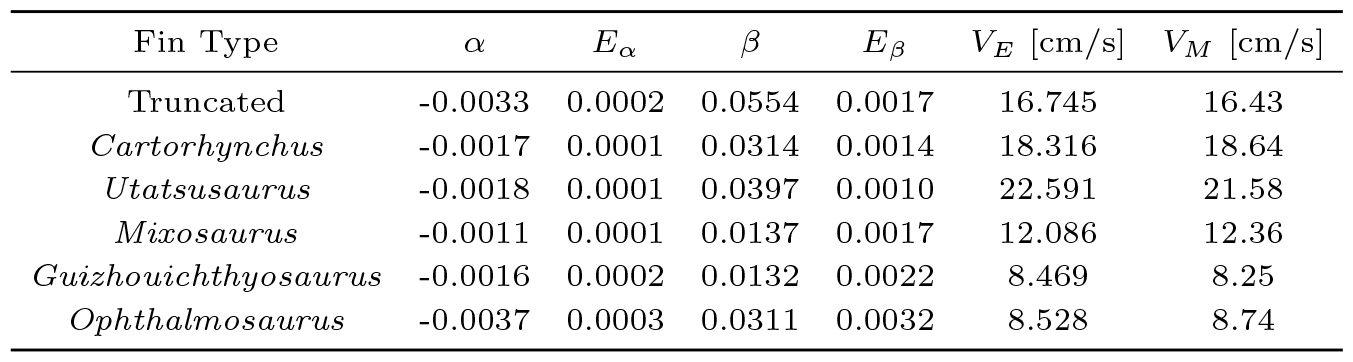
Self-propelled speed of the various fins. *α* and *E*_*α*_ are, respectively, the linear regression slope coefficient and its standard error; *β* and *E*_*β*_ are, respectively, the linear regression intercept coefficient and its standard error. *V*_*E*_ is the estimated self-propelled speed using linear regression, and *V*_*M*_ is its experimentally measured value. Poor performance from the last two fins is expected, as their thunniform morphology is more adapted to lift-based swimming, which the platform is not suited to provide in this study.

Pitch torque and forward thrust are not expected to be largely impacted by the change in flow speed, as drag is mostly applied in the direction of the flow. They should, however, change as the fins display a more distinct hypocercal morphology. Figure 7 shows that pitch is indeed mostly constant with a slight decline as the flow speed increases. Pitch values match the expected order with regards to the hypocercal fins, with the *Guizhouichthyosaurus, Utatsusaurus*, and *Mixosaurus* showing positive torque, while the other fins, all symmetrical, show values around zero.

Positive torque indicates that by having such hypocercal morphologies, those ichthyosauriforms had a tendency to passively plunge their snout towards the depths and thus may have helped compensate for positive buoyancy.

### Actuation Frequency Sweep

Figures 8, 9 and 10 show the average forward thrust (F_*x*_), vertical thrust (F_*z*_) and pitch torque (T_*y*_) during the frequency sweep experiment. Frequency is one of the main parameters that influences generated thrust, along with amplitude. At constant amplitude, frequency usually shows a linear relationship with thrust and swimming speed (Magnuson 1978). In our study, however, as we are not controlling it to remain constant, the tail swing amplitude tends to decrease as frequency increases, as shown in Figure 3 and 4. This phenomenon is related to the water resistance as well as the actuator and caudal fin stiffness, and it agrees with the results obtained by Wolf & Lauder (2021) in figure 8.4.B. As such, measured thrust can be expected to drop at high frequencies rather than progressing linearly. The combination of collected data and kinematic data reveals that longer tails often generate stronger forward propulsion. Once again, this is coherent with this experimental setup, but with proper stiffness, amplitude control, and fin-adapted motion, nature has shown that thunniform swimmers with high aspect ratio fins outperform the others in swimming speed.

**Fig. 3.**
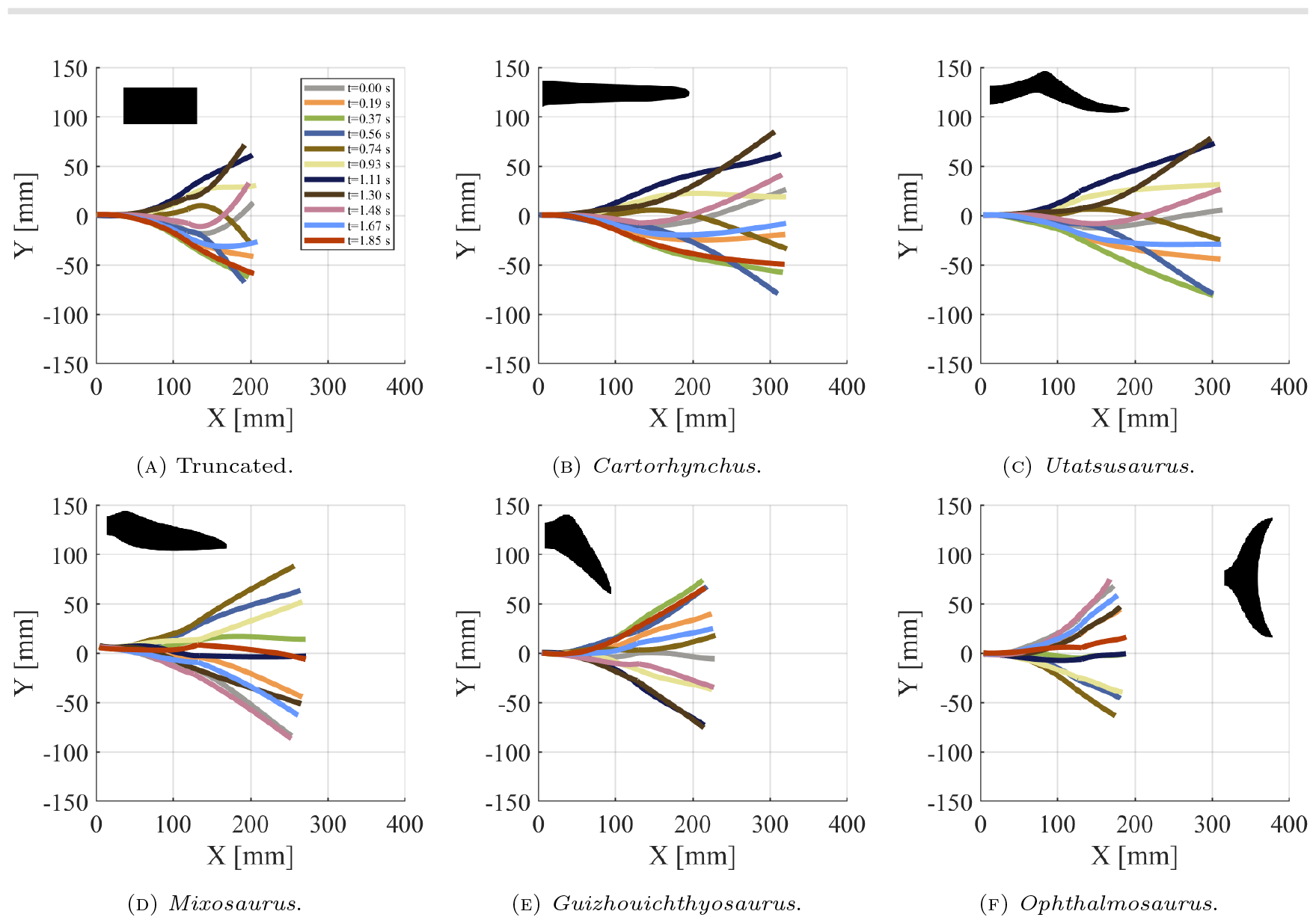
Midline kinematics of the truncated fin and five soft *Ichthyosauriformes* fins in the frequency sweep experiment. Oscillation amplitudes at undulation frequency of 0.6 Hz with constant flow speed of 5 cm/s.

**Fig. 4.**
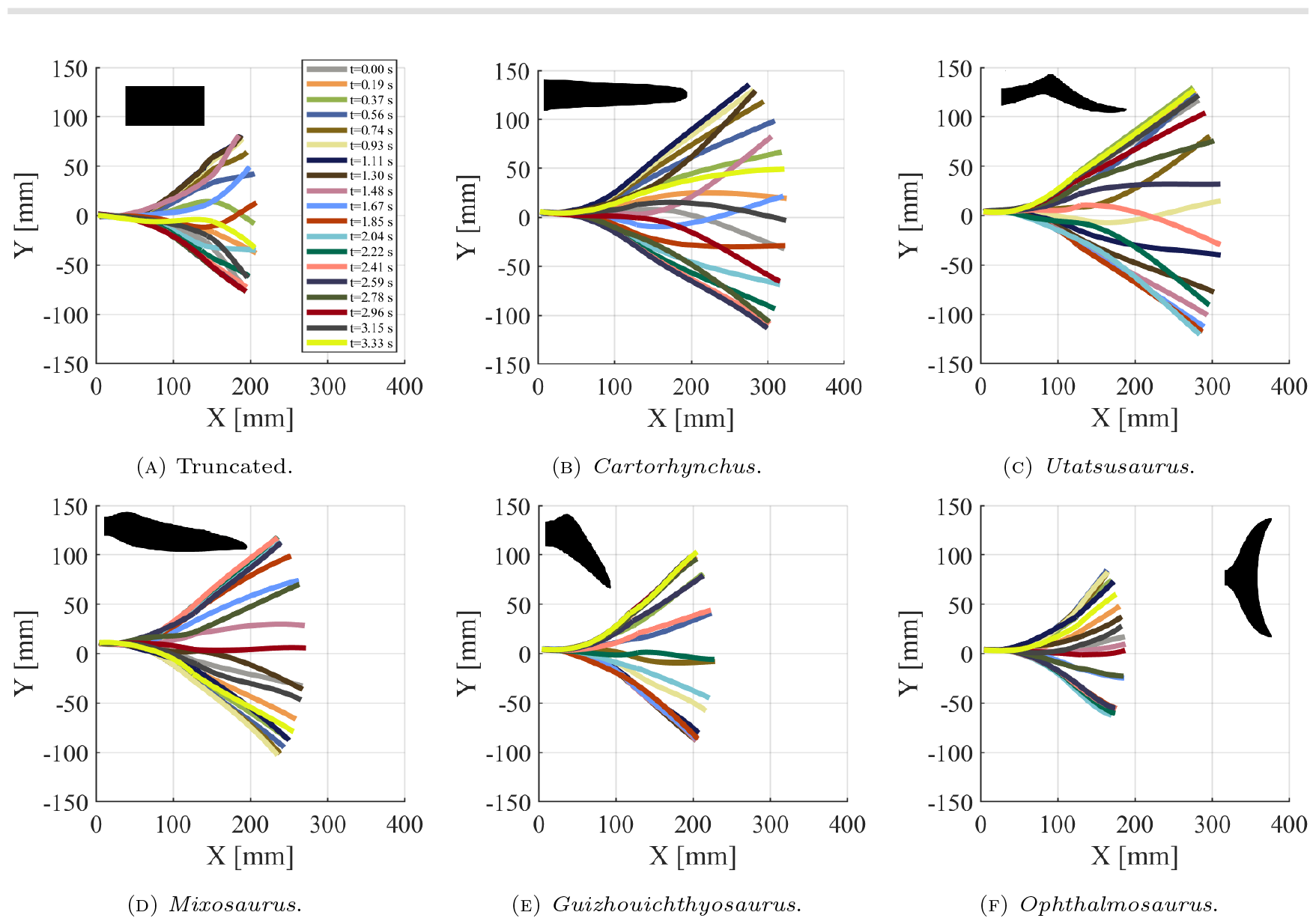
Midline kinematics of the truncated fin and five soft *Ichthyosauriformes* fins in the frequency sweep experiment. Oscillation amplitudes at undulation frequency of 0.3 Hz with constant flow speed of 5 cm/s.

### Limitation of Study

While selecting the caudal fin sections form for each model, different body parts have been represented and are therefore not homologous between all the chosen fins. For example, the tail of *Cartorhynchus* shown would include many more vertebrae compared to the *Mixosaurus* or the *Ophthalmosaurus*. As such, these tail depictions have different degrees of freedom and allow thus for different amounts of lateral bending.

The stiffness implementation and actuation of the platform do not align exactly with all the original specimens chosen for the study. Although it is suited to replicate subcarangiform or carangiform swimming, the longer forms (*Cartorhynchus, Utatsusaurus, Mixosaurus*, and potentially the longer *Guizhouich-thyosaurus* reconstruction) may have behaved more like anguilliform swimmers, with an overall lesser and more constant stiffness throughout the tail, allowing for more lateral freedom (Motani et al. 1996). The platform is also not suited to perform the oscillations associated with thunniform swimming (*Ophthalmosaurus*) appropriately (Sfakiotakis et al. 1999).

The stiffness of the swimming platform is not constant throughout its length; in fact, only the anterior part is actuated, while the posterior is a flexible passive foil. This particular mechanical structure allows the creation of different oscillating modes in the undulating tail, as can be seen in Figure 3 and Figure 4, similar to the results obtained by Wolf & Lauder (2021) in figure 8.3.C. The presence of oscillating modes in Figure 3 and Figure 4 alters the thrust generation while the robotic platform is actuated, as already hypothesized by Jusufi et al. (2017) in Figure 5, where a peak in thrust generation appears with an oscillating frequency of 0.55 Hz. The analysis of such modes is out of the scope of this paper, and it is left for future works.

## Discussion

Figures 5 and 8 give the reader a better understanding of how the generated thrust force changes with changes in flow speed and actuation frequency. The authors attribute the non-linear variation of the thrust force *F*_*x*_ in Figure 8 to the interaction among tail shape, frequency of oscillation, and flow speed (Liao et al. 2003). The increase in the flow speed (Figure 5) highlights the reduction of the forward thrust force generated while swimming upstream, with the *Oph-thalmosaurus* and *Guizhouichthyosaurus* fins being unable to “move forward” in such a flow with the current experimental setup.

**Fig. 5.**
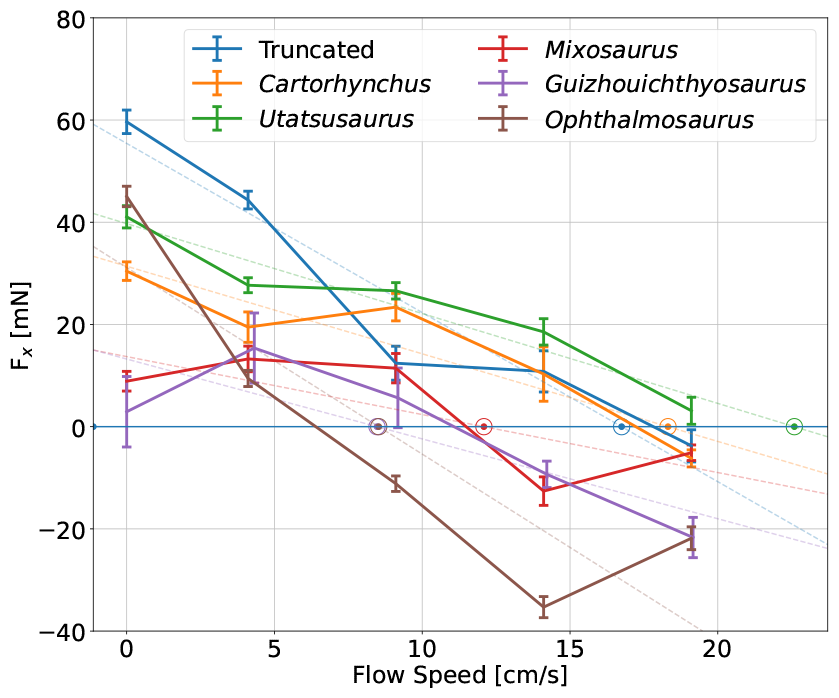
Forward thrust in the flow speed sweep experiment at a fixed actuation frequency of 0.55 Hz. The lines represent the mean value of the forward thrust over 10 undulation periods, while the error bar is the standard deviation of the mean. Dashed lines show the linear regression results, with the estimated self-propelled speeds highlighted by circles.

The results presented in the section above support the hypothesis previously proposed by Giammona (2021), Lauder (2000), and McGowan (1992) that heterocercal tails contribute to the balance of vertical forces caused by the fish buoyancy as they swim. The material properties, morphology, and swimming dynamics of fish tails are associated with their ecological niches (Naughton et al. 2021). We speculate that the symmetry or asymmetry of tail shapes may correlate with the need for additional pitch torque to maintain swimming stability within their ecological roles. Figures 6, 7, 9 and 10 show the effect of the presence of asymmetrical fins on the vertical force and pitch torque generation, validating the hypothesis that swimming mechanics combined with the tail shape can compensate for the animal’s positive or negative buoyancy. The *Guizhouichthyo-saurus* fin model, which presented the most extreme asymmetrical fin shape, as shown in Figure 1.D, was able to generate the greatest positive pitch torque *τ*_*y*_ while swimming among the tested fins. Figure 1.D suggests that in potentially positively buoyant swimmers such as *Guizhouichthyosaurus, Ophthalmosaurus* and *Mixosaurus*, swinging the heterocercal tail fin pushed down the animal’s snout and mitigated to an extent the upward force generated by the buoyancy, potentially allowing them to swim horizontally while cruising. On the other hand, the symmetric tail of the *Cartorhynchus* and the *Ophthalmosaurus* (see Figure 1.A and Figure 1.E respectively), as expected, did not generate significant pitch torque T_*y*_ while swimming, indicating a strict correlation between neutral buoyancy and symmetrical fin shape.

**Fig. 6.**
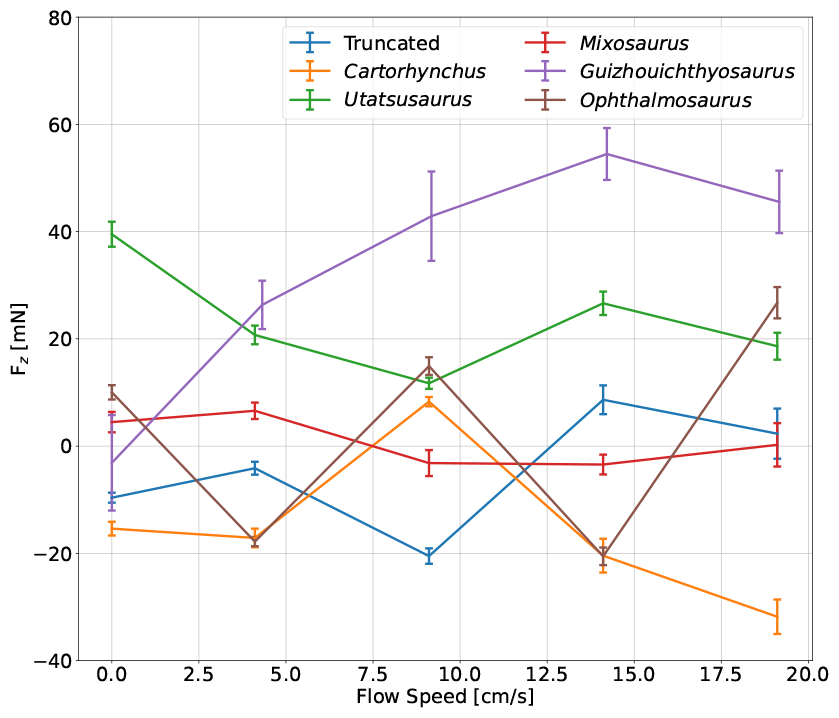
Vertical thrust in the flow speed sweep experiment at a fixed actuation frequency of 0.55 Hz. The lines represent the mean value of the forward thrust over 10 undulation periods, while the error bar is the standard deviation of the mean.

**Fig. 7.**
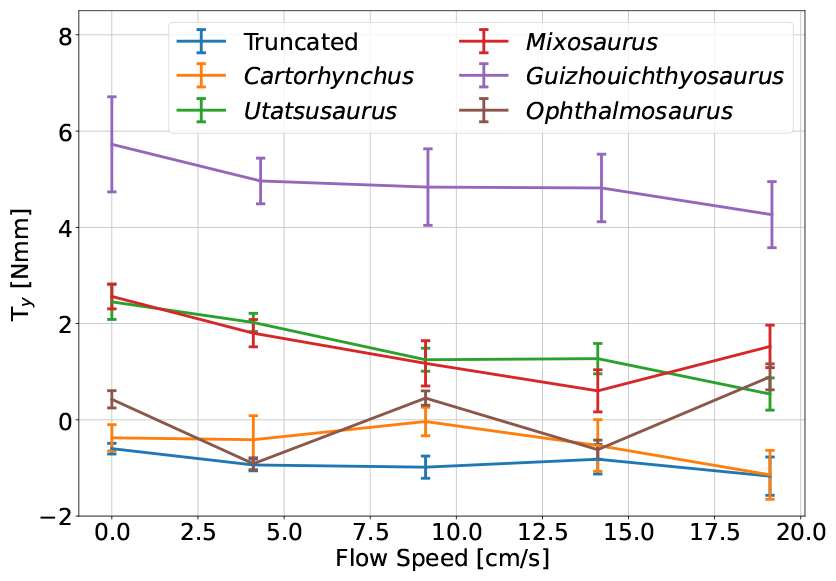
Pitch torque in the flow speed sweep experiment at a fixed actuation frequency of 0.55 Hz. The lines represent the mean value of the pitch torque over 10 undulation periods, while the error bar is the standard deviation of the mean.

**Fig. 8.**
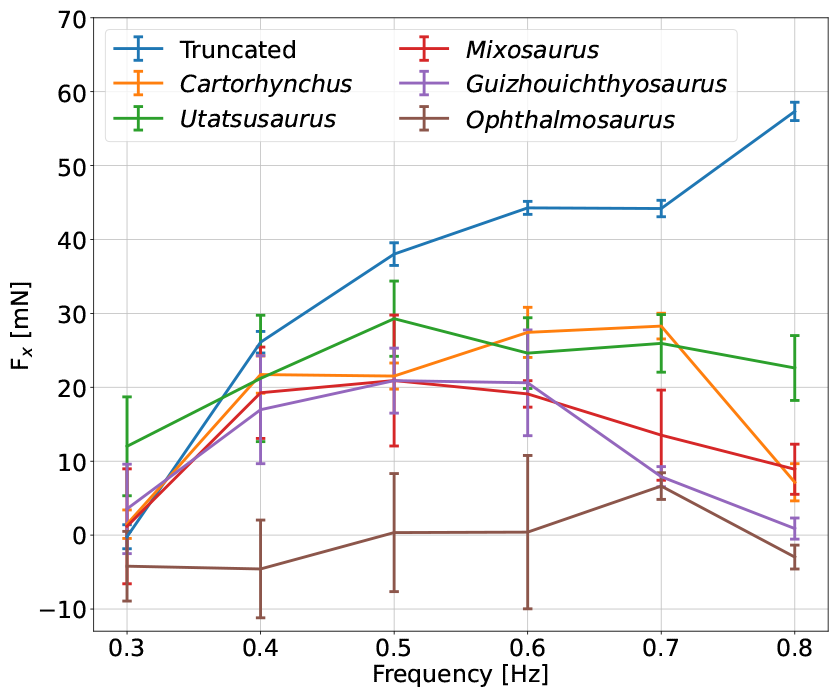
Forward thrust in the frequency sweep experiments at a fixed flow speed of 5 cm/s. The lines represent the mean value of the forward thrust over 10 undulation periods, while the error bar is the standard deviation of the mean.

**Fig. 9.**
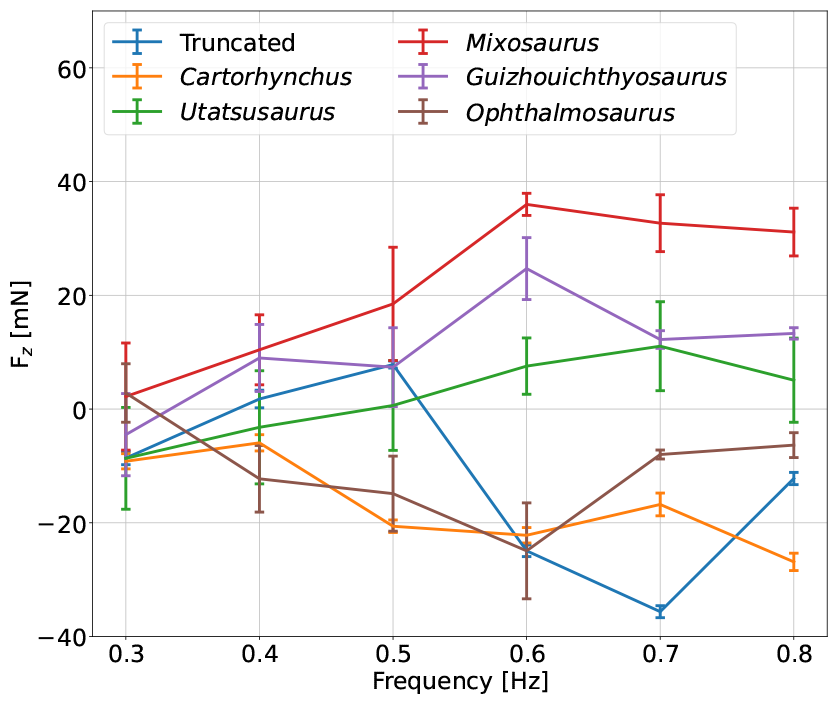
Vertical thrust in the frequency sweep experiments at a fixed flow speed of 5 cm/s. The lines represent the mean value of the forward thrust over 10 undulation periods, while the error bar is the standard deviation of the mean.

**Fig. 10.**
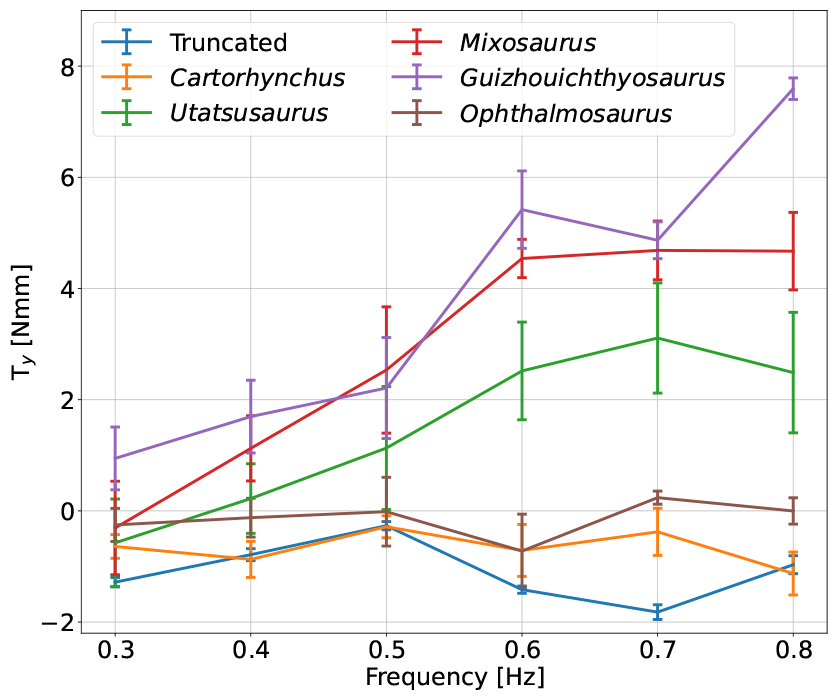
Pitch torque in the frequency sweep experiment at a fixed flow speed of 5 cm/s. The lines represent the mean value of the pitch torque over 10 undulation periods, while the error bar is the standard deviation of the mean.

## Conclusions

This work presented a novel application of a soft robot to evaluate the behavior of swimming animals without the need for severing or killing live specimens (McHenry et al. 1995, Long Jr 1998). The physical experiments performed with the robotic fish-shaped marine reptile allow us to understand better the swimming capabilities and characteristics of extinct animals and, more specifically, how the shape of the caudal fin and the animal’s buoyancy are related.

The outcome of this research is twofold. On the one hand, it shows that different types of fins lead to different pitch torques and that for heterocercal fins, the torques have the right sign for correcting buoyancy. On the other, it demonstrates the usefulness of a soft actuated robot as a tool to validate behaviors of extinct animals in real-life experiments, otherwise achievable only through computer simulations.

### Future Directions

The directions of future works are various but mainly focus on adapting the frequency oscillation of the tail to the flow speed to increase forward thrust production while reducing energy consumption. Another direction may include the analysis of the oscillating modes of the various fins as in (Tytell et al. 2016). An additional research topic may include the analysis of how the variation of the body stiffness while swimming, which influences the resistance to bending (Long Jr 1998), affects forward thrust generation.

## Acknowledgments

The authors thank Angelo Scioscia and Terence Fontana for valuable insights and Feiko Miedema for discussions on ichthyosaur anatomy.

